# *In vitro* activity of a novel antifungal compound, MYC-053, against clinically significant antifungal-resistant strains of *Candida glabrata, Candida auris, Cryptococcus neoformans*, and *Pneumocystis* spp

**DOI:** 10.1101/409896

**Authors:** G Tetz, M Collins, D Vikina, V Tetz

**Author notes:** To whom enquiries should be directed (G Tetz). Address correspondence to George Tetz.

## Abstract

An urgent need exists for new antifungal compounds to treat fungal infections in immunocompromised patients. The aim of the current study was to investigate the potency of a novel antifungal compound, MYC-053, against the emerging yeast and yeast-like pathogens *Candida glabrata, Candida auris, Cryptococcus neoformans*, and *Pneumocystis* spp. MYC-053 was equally effective against the susceptible control strains, clinical isolates, and resistant strains, with the minimum inhibitory concentrations (MIC) of 0.125–4.0 μg/mL. Notably, unlike other antifungal compounds, MYC-053 was effective against *Pneumocystis* isolates. MYC-053 was highly effective against preformed 48-h-old yeast biofilms, with the minimal biofilm eradication concentrations equal to 1–4 times MIC. The compound was not cytotoxic against L2 and A549 cell lines at concentrations over 100 μg/ml. Further, it possessed no apparent hemolytic activity up to 1000 μg/ml (the highest concentration tested). Overall, these data indicated that MYC-053 has a broad therapeutic window and may be developed into a promising antifungal agent for the treatment and prevention of invasive fungal infections caused by yeasts and yeast-like fungi in neutropenic patients.

---

In the last decade, invasive fungal infections caused by non-*albicans Candida* species and other less-common emerging yeasts, such as *Cryptococcus*, have become the leading cause of mortality in immunocompromised individuals (1–4).

*Candida glabrata* has emerged as the most common non-*albicans Candida* species and a causative agent of these invasive infections. It predominantly affects neutropenic patients, e.g., hematopoietic stem cell-transplant recipients, patients with HIV and diabetes, with high mortality rates (5). Systemic infections caused by these fungi include the associated cases of invasive candidiasis. Currently, non-*albicans* candidaemia is one of the most common causes of hospital-acquired bloodstream infections in the United States (6–9).

The main reason for the low efficacy of existing therapeutic options is the spread of fungal isolates with reduced susceptibility to the major classes of antimycotics (polyenes, azoles, and echinocandins) (10). The spread of multidrug resistant strains of *C. glabrata* in the US, displaying resistance to at least two classes of antifungal drugs, severely limiting treatment options, and consistently associated with increased mortality was described in recent studies (11–13). Notably, resistance to echinocandins, a novel class of antifungal agents, is also increasing among *C. glabrata* isolates, with the reported resistance rate of 3–12% in different countries (14,15).

Another global healthcare concern is the emerging multidrug-resistant pathogenic *Candida auris* (16,17). Unlike most other *Candida* spp., this fungus is commonly transmitted within health care facilities (18). One of the reasons for the nosocomial spread of *C. auris* is its survival on surfaces in healthcare facilities for many days (19). Recently, *C. auris* infections have been reported to cause outbreaks in over 50 healthcare facilities in 15 countries (20). Moreover, drug resistance of *C. auris* exceeds that of *C. glabrata*, with over 41% isolates reportedly resistant to at least two antifungal classes (18).

*Cryptococcus neoformans* is another opportunistic pathogen and an etiologic agent of cryptococcosis, a life-threatening infection in immunocompromised hosts, particularly HIV-infected, with cancer, or solid-organ transplant recipients (4). Although the rates of cryptococcosis have dropped substantially since the development of highly active antiretroviral therapy, the mortality of HIV patients associated with cryptococcal meningitis remains high. One of the causes of treatment failure is the emergence of azole-resistant and heteroresistant mutants (21–23).

Another important yeast-like fungal pathogens of the immunocompromised neutropenic host are *Pneumocystis* species, which cause pneumocystis pneumonia (PcP) (24). According to the statistics of the Centres for Disease Control and Prevention, the incidence of PcP among hospitalized HIV patients in the US is 9% (CDC. 2014. Pneumocystis pneumonia statistics. http://www.cdcgov/fungal/diseases/pneumocystis-pneumonia/statisticshtml). *Pneumocystis* spp., originally classified as protozoa, are now classified as fungi but are not susceptible to anti-fungal drugs (REF). Hence, the main therapeutic options of preventing and treating PcP are a combination of trimethoprim, sulfamethoxazole, and aerosolized pentamidine (25). Nevertheless, the therapy is ineffective for many AIDS patients, with the current mortality rate of PcP remaining high and ranging from 5 to 40%, while the mortality is 100% in non-treated patients (26).

In addition to the spread of antifungal resistance, biofilm formation is a global concern associated with all fungal infections, which leads to therapy failure (27–29). Biofilms formed during invasive fungal infections by yeasts and yeast-like fungi retard the penetration and diffusion of antifungal agents and enabling the fungi to survive in the presence of high antibiotic concentrations (30,31). Therefore,development of novel antifungal agents that would be effective against resistant fungi that are prevalent among neutropenic patients is at the forefront of today’s medicine. In the current study, the fungicidal activity of a novel antifungal compound, MYC-053 [sodium 5-[1-(3,5-dichloro-2-hydroxyphenyl)methylideneamino]-6-methyl-1,2,3,4-tetrahydro-2,4-pyrimidinedionate] (Fig. 1), which is not related to any existing classes of antifungal agents, was investigated against planktonic and biofilm-forming *Candida* spp., *Cryptococcus* spp., and *Pneumocystis* spp.

**FIG 1.**
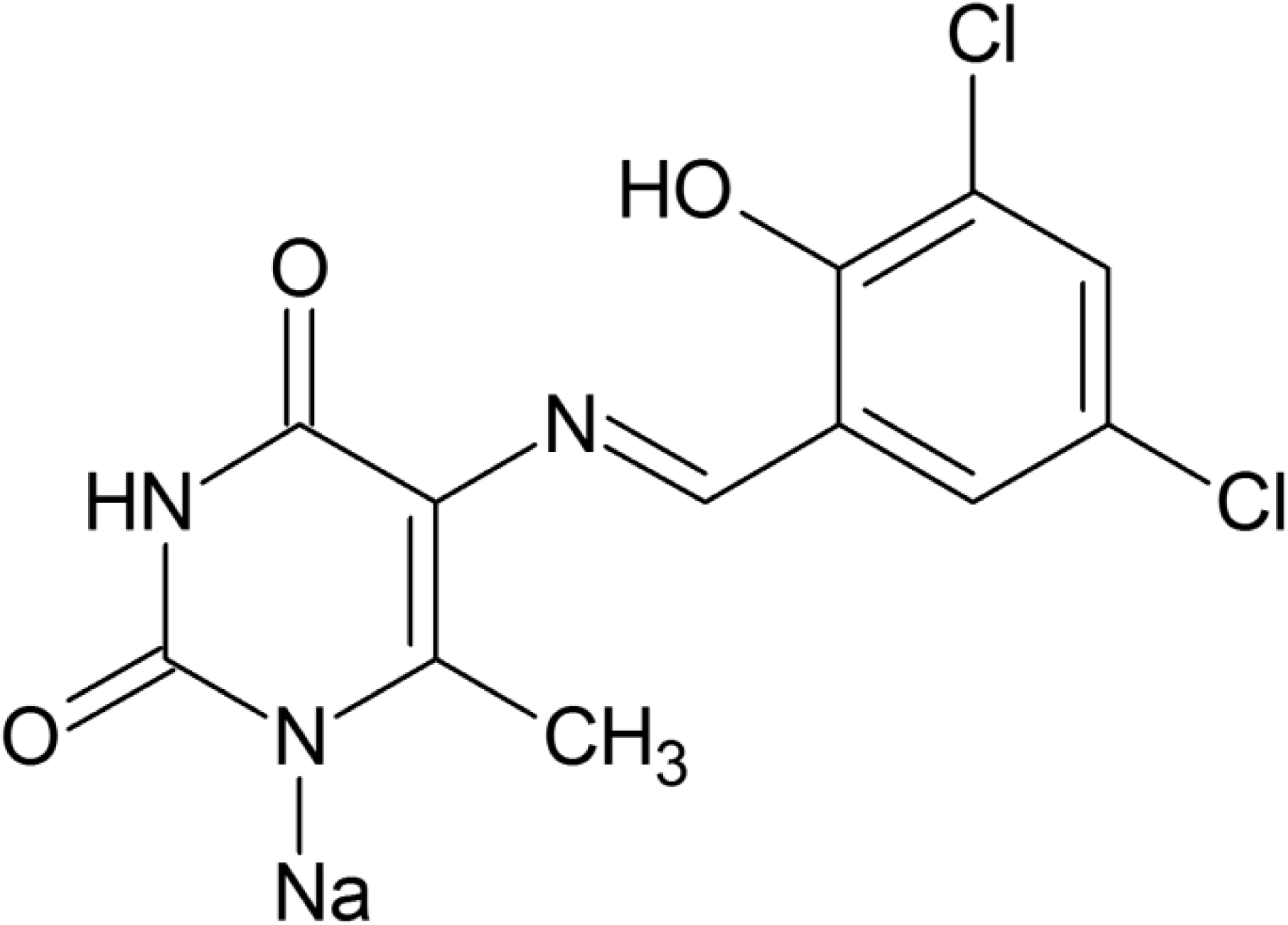
Chemical structure of MYC-053.

**FIG 2.**
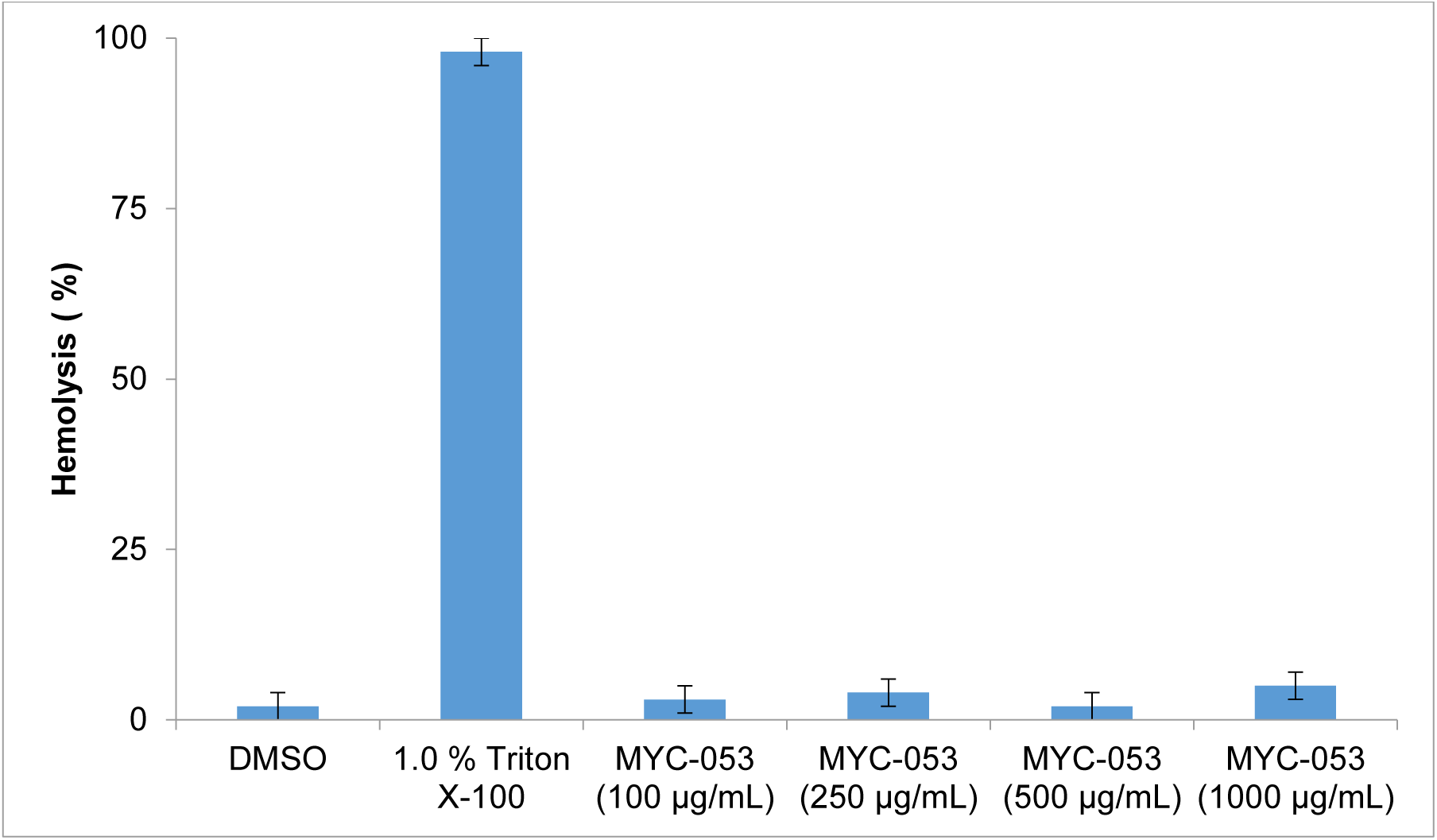
Percent hemolysis of human erythrocytes exposed to 100–1000 μg/ml of MYC-053. Positive control (Triton X-100); negative control (DMSO). No hemolysis was observed with any of the tested concentrations of MYC-053 up to 1000 μg/ml. Results represent the means from 3 experiments which each contained three technical replicates.

## RESULTS

***In vitro* antifungal activity of MYC-053 against *C. glabrata, C. auris*, and *C. neoformans*.** The efficacy of MYC-053 against a panel of 20 *C. glabrata* strains [including seven flcconazole (FLC)-resistant and four caspofungin (CAS)-resistant strains], five *C. auris* strains, and 18 *C. neoformans* strains (including 12 FLC-resistant strains)was determined by the broth microdilution method (Table 1). Minimum inhibitory concentrations 50 (MIC_50_) values of MYC-053 for *C. glabrata* strains varied from 0.125 to 0.5 μg/mL, while Minimum inhibitory concentrations 100 (MIC_100_) values were in the low μg/mL range (1–4 μg/ml). *C. auris* and *C. neoformans* strains were also sensitive to this compound, with MIC_50_ and MIC_100_ of 0.5–4 μg/ml. Notably, the antifungal activity of MYC-053 against the susceptible strains, including the control C. *glabrata* ATCC 90030 and C. *neoformans* ATCC 90112 strains, was a lower than that of FLC but higher than that one of CAS. In contrast, while certain resistant clinical isolates exhibited reduced susceptibility to FLC and CAS, they were highly sensitive to the same concentrations of MYC-053 as the control strains (Table 1).

**TABLE 1.**
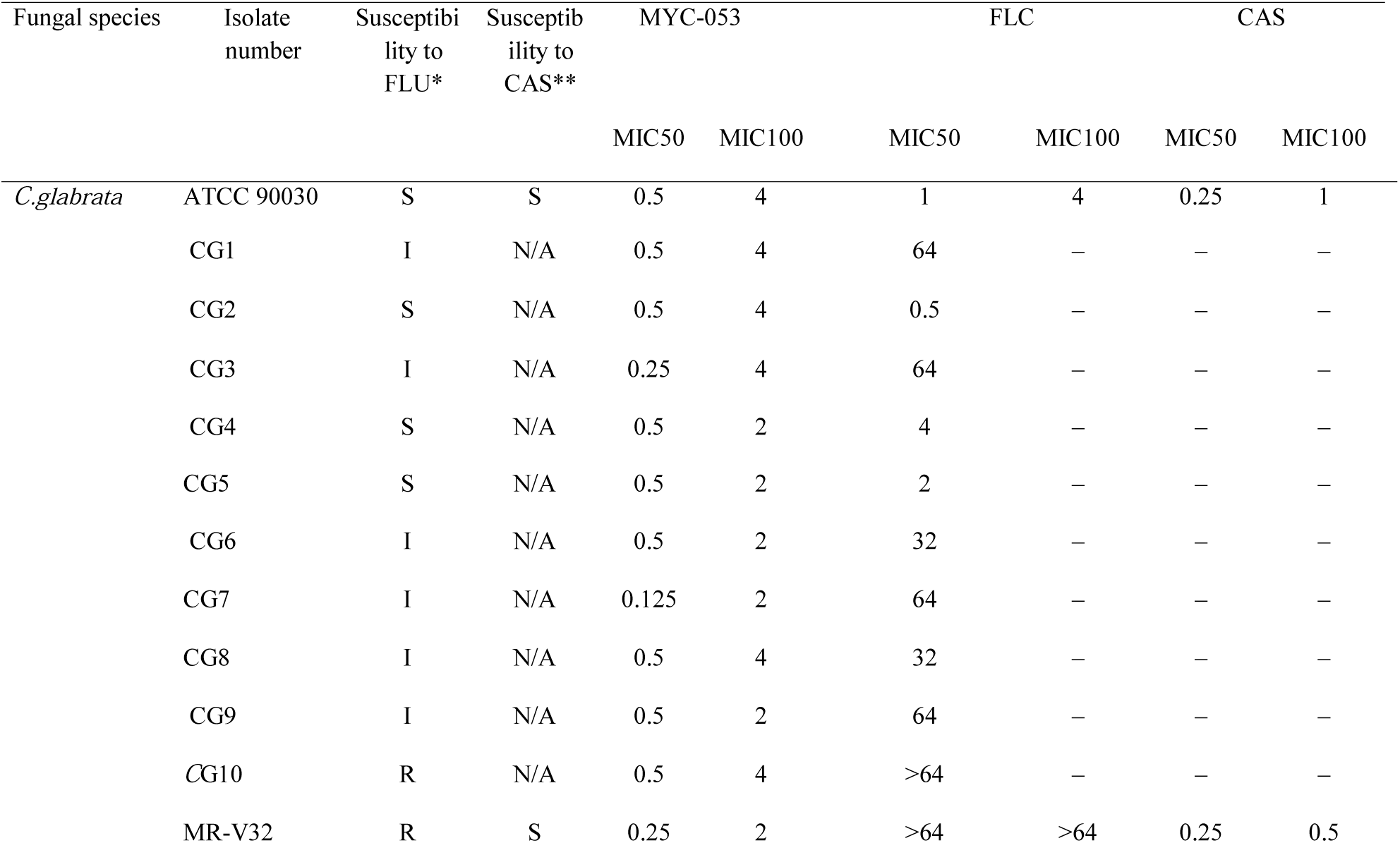
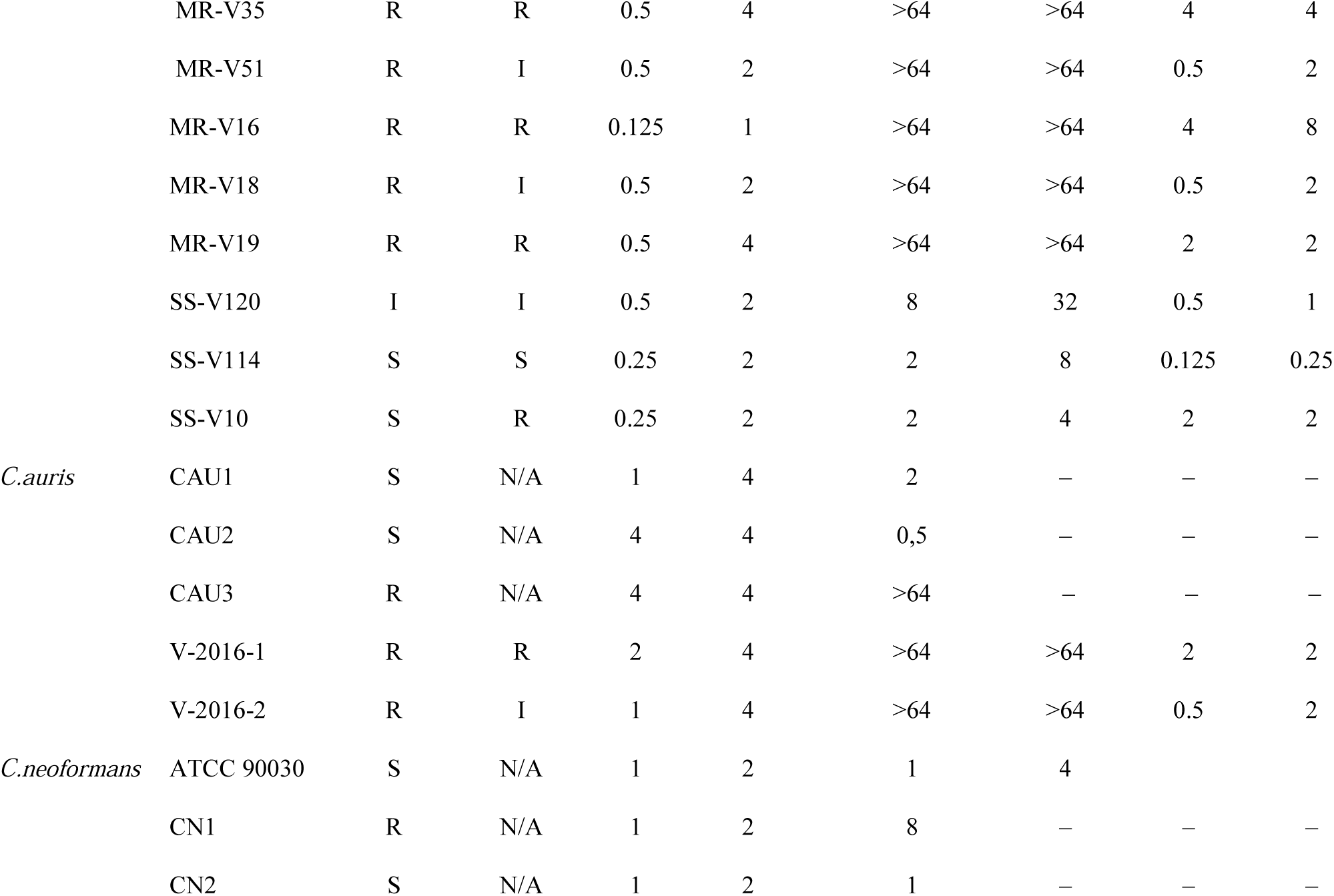
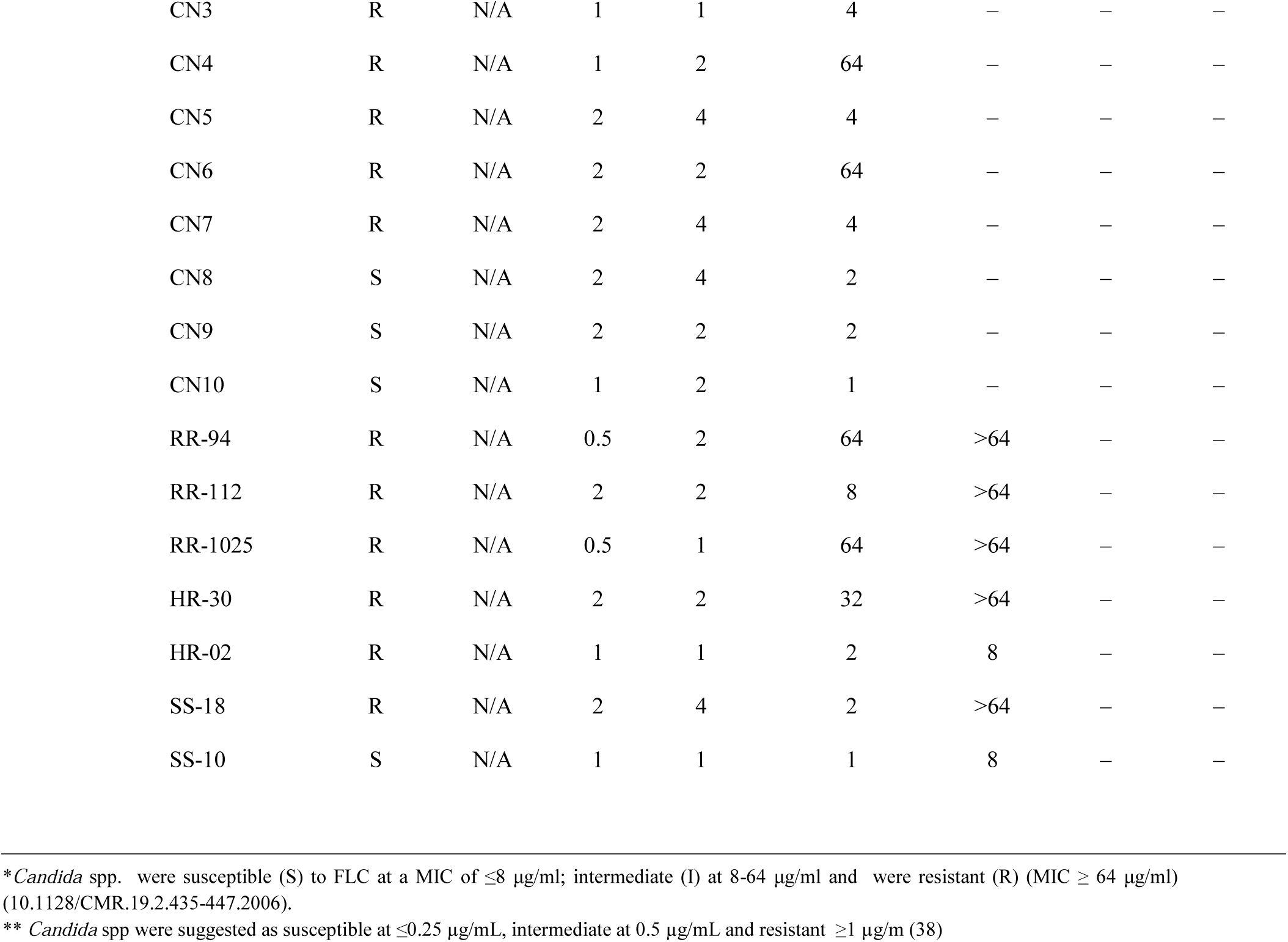
MIC of MYC-053 and other antifungal agents against *C. glabrata* and *C. auris*

### *In vitro* antifungal activity of MYC-053 against *Pneumocystis carinii* and *Pneumocystis murina*

The responses of *P. carinii* and *P. murina* to MYC-053 were evaluated by a cytotoxicity assay based on ATP-driven bioluminescence (32). The results, expressed as IC_50_ after 24, 48, and 72 hours of exposure to the drug, were assigned activity ranks based on the degree of reduction of ATP compared to untreated controls (33,34) (Table 2). The exposure of *P. carinii* to 1 μg/ml MYC-053 for 72 h resulted in a level of ATP reduction that was slightly lower than that of pentamidine and can be considered as moderate activity. However, the increase of MYC-053 concentration to 10 μg/ml resulted in lower of ATP pools, compared with 1 μg/ml pentamidine. The inhibitory effect of MYC-053 against *P. murina* was higher than against *P. carinii*. In this assay, MYC-053 demonstrated comparable activity against *P. murina* as pentamidine, with over 94.4% reduction of the ATP pool following 72-h exposure, which is considered to indicate marked activity on the efficacy scale (34). Overall, MYC-053 effectively reduced the ATP content of both *Pneumocistis* species at microgram levels. The following IC_50_ values were calculated over 3 d of *P. carinii* exposure to MYC-053: 3.90 μg/ml at 24 h; 2.56 μg/ml at 48 h; and 1.61 μg/ml at 72 h. Against *P. murina*, the IC_50_ values were: 3.30 μg/ml at 24h; 1.50 μg/ml at 48 h; and 0.165 μg/ml at 72 h.

**TABLE 2.** IC_50_ values for MYC-053 for *P. carinii* and *P. murina* following different exposure times in the ATP assay. Results represent the means from 3 experiments which each contained three technical replicates.

| Drug | 24h | 48h | 72h |
| --- | --- | --- | --- |
| *P.carinii* |  |  |  |
| Ampicillin 10$\mu\text{g/ml}$ | 7.84 | 1.51 | 0 |
| Pentamidine 1$\mu\text{g/ml}$ | 81.14 | 86.58 | 86.57 |
| MYC-053 50$\mu\text{g/ml}$ | 96.81 | 97.61 | 99.21 |
| MYC-053 10$\mu\text{g/ml}$ | 68.26 | 90.29 | 95.58 |
| MYC-053 1$\mu\text{g/ml}$ | 14.95 | 11.08 | 26.77 |
| MYC-053 0.1$\mu\text{g/ml}$ | 0 | 3.20 | 13.42 |
| $\text{IC}_{50}$ | 3.90 ± 2.0$\mu\text{g/ml}$ | 2.56 ± 0.57$\mu\text{g/ml}$ | 1.61 ± 1.72$\mu\text{g/ml}$ |
| *P. murina* |  |  |  |
| Ampicillin 10$\mu\text{g/ml}$ | 2.86 | 0.26 | 0 |
| Pentamidine 1$\mu\text{g/ml}$ | 92.07 | 97.70 | 98.12 |
| MYC-053 50$\mu\text{g/ml}$ | 97.84 | 98.89 | 98.77 |
| MYC-053 10$\mu\text{g/ml}$ | 76.56 | 98.51 | 97.92 |
| MYC-053 1µg/ml | 1.082 | 42.11 | 94.42 |
| MYC-053 0.1µg/ml | 0 | 0 | 27.82 |
| IC50 | 3.30 +/-0.19µg/ml | 1.50 +/-0.13µg/ml | 0.165 +/-0.06µg/ml |

### Activity of MYC-053 against *C. glabrata* and *C. neoformans* biofilms

The anti-biofilm effect of MYC-053, FLC, and CAS on preformed 48-h-old *C.glabrata* biofilms, and the effect of MYC-053 and FLC on cryptococcal biofilms were evaluated (Table 3). Preformed biofilms were exposed to drugs provided at concentrations equal to 1–64 times those of their MICs. MYC-053 significantly reduced the CFU of preformed biofilms of both *C. glabrata* and *C. neoformans* after 24 h of incubation, starting at a concentration of 1× MIC. MYC-053 at a concentration of 1× MIC decreased the number of viable fungi in all strains by more than 50%; this value was recorded as the minimum biofilm eradication concentration, MBEC_50_. Moreover, MYC-053 was the only drug that showed MBEC_90_ values equal to 1– 4 times its MIC. In the assay, higher relative concentrations of FLC and CAS were required to kill yeasts in preformed biofilms than MYC-053. The MBEC_50_ and MBEC_90_ values of FLC and CAS against the tested preformed *C. glabrata* biofilms were equal to 4–64 times and 1–32 times their MICs, respectively. Similar data with high relative MBEC_50_ and MBEC_90_ concentrations of FLC required were obtained against *C. neoformans* biofilms. CAS efficacy was not tested against *C. neoformans* biofilms in this assay, as this microorganism is known to be resistant both *in vitro* and *in vivo* to echinocandins (REF).

**TABLE 3.** Susceptibility of 48-h-old *C. glabrata* biofilms to MYC-053, FLC, and CAS, expressed as multiples of MIC values. Results represent the means from 3 experiments which each contained three technical replicates.

| Fungal species | Isolate number | MYC-053 |  | FLC |  | CAS |  |
| --- | --- | --- | --- | --- | --- | --- | --- |
|  |  | MBEC<sub>50</sub> | MBEC<sub>90</sub> | MBEC<sub>50</sub> | MBEC<sub>90</sub> | MBEC<sub>50</sub> | MBEC<sub>90</sub> |
| *C.glabrata* | ATCC 90030 | 1 | 2 | 4 | 4 | 1 | 4 |
|  | MR-V32 | 1 | 2 | 4 | >64 | 4 | 16 |
|  | MR-V35 | 1 | 1 | 8 | >64 | 8 | 4 |
|  | MR-V51 | 1 | 2 | 4 | >64 | 2 | 2 |
|  | MR-V16 | 1 | 2 | 4 | >64 | 4 | 32 |
|  | MR-V18 | 1 | 4 | 32 | >64 | 2 | 16 |
|  | MR-V19 | 1 | 2 | 16 | 32 | 4 | 32 |
|  | SS-V120 | 1 | 1 | 8 | 8 | 2 | 4 |
|  | SS-V114 | 1 | 4 | 4 | 32 | 4 | 32 |
|  | SS-V10 | 1 | 1 | 4 | 16 | 16 | 32 |
| *C.neoformans* | ATCC 90030 | 1 | 2 | 2 | 16 | – | – |
|  | RR-94 | 1 | 4 | 4 | >64 | – | – |
|  | RR-112 | 1 | 4 | 32 | >64 | – | – |
|  | RR-1025 | 1 | 1 | 8 | 32 | – | – |
|  | HR-30 | 1 | 1 | 16 | 64 | – | – |
|  | HR-02 | 1 | 2 | 64 | >64 | – | – |
|  | SS-18 | 1 | 2 | 8 | 16 | – | – |
|  | SS-10 | 1 | 1 | 4 | 16 | – | – |

### MYC-053 possesses no apparent cytotoxic or hemolytic activities

Since the *in vitro* antifungal activity of MYC-053 was demonstrated in different experimental set-ups, we next evaluated the possible cytotoxicity of this molecule in two eukaryotic cell lines. L2 and A549 cells were treated with increasing concentrations of MYC-053 (0.1–100 μg/ml) for up to 72 h. The cytotoxicity was evaluated using an ATP release assay, as described in the Materials and methods with three independent repeats at each time point (35). According to the EU classification criteria, MYC-053 was not toxic to either cell line, with IC_50_ values over 100 μg/ml (36,37). To evaluate the possible hemolytic activity of MYC-053, human erythrocytes were exposed to increasing concentrations of the compound (up to 1000 μg/ml) for 6 h. No significant difference was observed between the control (dimethyl sulfoxide, DMSO) and MYC-053.

## DISCUSSION

In the United States, invasive candidiasis has become a common healthcare-associated infection. Currently, non-*albicans Candida* spp. are the major cause of morbidity in immunocompromised patients, including patients with HIV, cancer, hematological malignancies, diabetes, and solid organ transplants. The prevalence of *C. glabrata* among non-*albicans Candida* infections has been constantly increasing over time, and this fungus is now registered as the most frequently isolated fungus species from patients (38).The main reasons for the failure of the existing therapeutic options is an increase in antifungal resistance and low sensitivity of fungal biofilms to the existing antibiotics (39–41). Another serious global health threat represented by non-*albicans Candida* species is *C. auris*, a recently discovered, rapidly emerging fungal pathogen (42). Infections due to *C. auris* are predominantly hospital-acquired and up to 30% are represented by multidrug resistant strains, including ones resistant to all three classes of antifungal agents, and are associated with high mortality rates (43). Although the incidence of infections caused by *Cryptococcus* spp. and *Pneumocystis* spp. has decreased since the introduction of HAART, which partially restores the function of the immune system, opportunistic infections caused by these fungi continue to contribute to the mortality of the immunocompromised hosts (44,45).

In the current study, we described a novel antifungal drug candidate, MYC-053, which exhibited a high level of antimicrobial activity against *C. glabrata, C. auris, C. neoformans*, and *Pneumocistis* spp. *in vitro*. Importantly, the MIC experiment revealed that MYC-053 exerted a pronounced cidal effect against resistant fungal isolates at concentrations identical to the ones killing susceptible control fungal strains. These data correspond well with the notion that MYC-053 is a representative of a novel chemical class of antifungal agents; it is not relevant to the existing antifungal agents whose use is frequently characterized by cross-resistance (46). Notably, MYC-053 was effective against *C. auris* that is often multidrug-resistant (47). Although we have only tested the activity of MYC-053 against five *C. auris* strains, low SEM values in the assay suggested high precision of the measurements allowing us to determine the mean MIC_50_ as 1 μg/ml and MIC_100_ as 4 μg/ml. Despite the fact that MIC values of MYC-053 against *C. auris* were higher than against *C. glabrata*, these values were nonetheless promising given the low susceptibility of certain tested strains to FLC and CAS, with MIC values over 64 μg/mL for these antifungals.

MYC-053 was also effective against *C. neoformans*, with MIC values starting at 1.0 μg/ml. These concentrations were dramatically different from the FLC MIC values. Although we did not test the sensitivity of *C. neoformas* strains against other azoles, it is known that this fungus is commonly cross-resistant to other antifungal agents of this class, including voriconazole (48,49). Therefore, we propose that MYC-053 might be effective against other azole-resistant strains of C. *neoformans*.

This investigation also revealed that MYC-053 was effective against *Pneumocystis* spp., other yeast-like pathogens that are challenging to treat in neutropenic patients. The anti-pneumocystis activity of MYC-053 was promising since, despite being originally classed as protozoa, *Pneumocystis* spp. are now classified as fungi and continue to be generally treated with antibacterial and antiprotozoan medications (32, 50). The determination of ATP levels for the assessment of MYC-053 activity against *Pneumocystis* constitutes a highly sensitive assay enabling the reduction of the number of tested organisms (33). The activity of MYC-053 was considered as marked and was comparable to the activity of pentamidine against *P. murina* at the 72-h time point. To the best of our knowledge, MYC-053 is the first new synthetic compound that can be potentially used against *Pneumocistis* spp., *Candida* spp., and *Cryptococcus* spp…

To determine whether MYC-053 was active against preformed yeast biofilms, its MBEC values were evaluated at concentrations equal to multiples of the MIC values. In the assay, 48-h-old preformed biofilms of susceptible and clinical isolates were evaluated. At a concentration equal to MIC, MYC-053 caused a 50% reduction of the viable cell counts in all studied fungal biofilms. The MBEC_90_ values of MYC-053 were equal to 1–4-time multiples of MIC values. Notably, the MIC/MBEC_50/90_ ratios of MYC-053 were significantly lower than those of the control antifungals FLC and CAS. In summary, MYC-053 was equally effective against sessile and planctonic non-resistant organisms and multiresistant clinical isolates.

Importantly, the potent antifungal activity of MYC-053 was not associated with any cytotoxicity and hemolytic activity, as evaluated in a 72-h assay, suggesting a high therapeutic index. Therefore, considering the clinical efficacy of an antifungal agent based on its anti-biofilm activity but not on the MIC criterion, it would be easy to conclude that the concentrations of FLC and CAS required to eliminate biofilms formed by the resistant strains of *C. glabrata* and *C. neoformans* cannot be achieved in human because of the toxicity limitations (51,52). At the same time, the high therapeutic window of MYC-053, with low toxicity and high anti-biofilm activity render the data promising. These findings suggest that MYC-053 may be used to treat of fungal infections that involve biofilm formation.

Taken together, the results of the current study on the efficacy of MYC-053 against certain yeasts and yeast-like pathogens, including ones in biofilm state, indicate the possibility of developing MYC-053 further into an antifungal drug candidate; however, it requires more *in vivo* research.

## MATERIALS AND METHODS

### The test substance and antimicrobials

MYC-053 was synthesized by TGV-inhalonix Inc. (Wilmington, DE); FLC, CAS, and pentamidine were purchased from Sigma Aldrich (St Louis, MO).

### Fungal strains

Forty-four fungal species were used in this study. *C. glabrata* CG1, *C. glabrata* CG2, *C. glabrata* CG3, *C. glabrata* CG4, *C. glabrata* CG5, *C. glabrata* CG6, *C. glabrata* CG7, *C. glabrata* CG8, *C. glabrata* CG9, *C. glabrata* CG10, *C. auris* CAU1, *C. auris* CAU2, *C. auris* CAU3, *C. neoformans* CN1, *C. neoformans* CN2, *C. neoformans* CN3, *C. neoformans* CN4, *C. neoformans* CN5, *C. neoformans* CN6, *C. neoformans* CN7, *C. neoformans* CN8, *C. neoformans* CN9, and *C. neoformans* CN10 were obtained from the Fungus Testing Laboratory at the University of Texas Health Science Center (San Antonio, TX). *C. glabrata* MR-V32, *C. glabrata* MR-V35, *C. glabrata* MR-V51, *C. glabrata* MR-V16, *C. glabrata* MR-V18, *C. glabrata* MR-V19, *C. glabrata* SS-V120, *C. glabrata* SS-V114, *C. glabrata* SS-V10, *C. auris* V-2016-1, *C. auris* V-2016-2, *C. neoformans* RR-94, *C. neoformans* RR-112, *C. neoformans* RR-1025, *C. neoformans* HR-30, *C. neoformans* HR-02, *C. neoformans* SS-18, and *C. neoformans* SS-10 were provided by Dr. V. Tetz (Human Microbiology Institute) from a private collection. *P. carinii* and *P*. *murina* were obtained from Melanie Cushion’s laboratory at the University of Cincinnati (Cincinnati, OH). The control strains were C. *glabrata* ATCC 90030 and *C. neoformans* ATCC 90112 obtained from the American Type Culture Collection (ATCC, Rockville, USA). *C. glabrata* and *C. auris* isolates were subcultured on Sabouraud dextrose agar before testing (Oxoid Ltd., Basingstoke, UK).

### *In vitro* antifungal susceptibility testing

Microdilution broth susceptibility testing was performed in duplicate according to the CLSI M27-A3 method in RPMI-1640 growth medium (Sigma Aldrich) to determine the MIC values (53). Standard inoculum for yeast testing was 2.5 × 10^3^ CFU/ml. FLC and CAS were dissolved in DMSO (Sigma Aldrich), whereas MYC-053 was dissolved in sterile water. MIC_50_ was defined as the lowest concentration of a drug that caused a 50% decrease in culture turbidity compared to the growth control. MIC_100_ was defined as the lowest concentration of the drug that resulted in no visual growth after 24 h of incubation at 35°C. Fungal isolates were categorized as susceptible, intermediate, or resistant, according to the susceptibility breakpoints for antifungals based on CLSI criteria (54,55). *C. glabrata* strains were susceptible to FLC at MIC ≤ 8 μg/ml; intermediate at 8–64 μg/ml; and resistant at MIC ≥ 64 μg/ml. MIC values for CAS were interpreted to indicate susceptible strains at ≤ 0.25 μg/ml; intermediate strains at 0.5 μg/ml; and resistant strains ≥1 μg/ml. No established *C. auris*-specific susceptibility breakpoints are currently available. Therefore, the same susceptibility breakpoints as for *C. glabrata* were used. *C. neoformans* strains were susceptible to FLC. No accepted *Cryptococcus* interpretive breakpoints are currently available. Therefore, only potential breakpoints for FLC against *C. neoformans* were used, as follows: susceptible, ≤ 2 mg/l; resistant, > 2 mg/l (56).

### *In vitro P. carinii* and *P. murina* ATP assays

MYC-053 was diluted directly in the culture medium (0.1, 1, 10, and 50 μg/ml). The culture medium was RPMI-1640 containing 20% horse serum, 1% Minimum essential medium (MEM) vitamin solution, 1% MEM Nonessential amino acids (NEAA), 200 U/ml penicillin, and 0.2 mg/ml streptomycin (Sigma Aldrich). The medium alone and medium containing 10 μg/ml ampicillin (Sigma Aldrich) were the negative controls. Medium supplemented with 1 μg/ml pentamidine isethionate was the positive control. Cryopreserved and characterized *P. carinii* isolated from the rat lung tissue and *P. murina* isolated from the mouse lung tissue were distributed into triplicate wells of 48-well plates (final volume of 500 μl and a final concentration of 5 × 10^7^ nuclei/ml for *P. carinii* and 5 × 10^6^ for *P. murina*). The controls and diluted compounds were added to the cultures and incubated at 35°C under 5% CO_2_. After 24, 48, and 72 h, 10% of the well volume was removed and ATP content was determined using the ATP-Lite luciferin-luciferase assay (Perkin-Elmer, Waltham, MA). The ATP-associated luminescence was determined using a spectrophotometer (PolarSTAR OPTIMA, BMG-Labtech, Germany). An each sample was examined microscopically on the final day of the assay to rule out the presence of bacteria. An quench control assay to determine compound interference in the luciferin/luciferase reaction was negative at all tested concentrations. Background luminescence was subtracted and triplicate well readings were averaged. For each time point, the percent reduction in ATP content in all groups was calculated as follows: [media control – (experimental/media control)] × 100. The 50% inhibitory concentration (IC_50_) was calculated using INSTAT linear regression program (GraphPad Software Inc., San Diego, CA).

### Effect of MYC-053 on preformed fungal biofilms

A standardized *C. glabrata* or *C. neoformans* culture inoculum (200 μl; 5 × 10^5^ CFU/ml) in RPMI-1640 was added to each well of a 96-well round-bottom polystyrene tissue culture microtiter plate (Sarstedt, Nümbrecht, Germany) (57,58). Following 48-h incubation at 35°C, biofilm samples were washed twice with phosphate-buffered saline to remove non-adherent cells and then exposed for 24 h to 200 μl of RPMI-1640 containing MYC-053, FLC, or CAS at concentrations equal to 1, 2, 4, 8, 16, 32, and 64 times their MICs. Untreated biofilms were used as negative controls. The number of viable fungi in the biofilm was determined by estimating the CFU number. Briefly, to estimate the CFU number, following exposure, well contents were aspirated to prevent antimicrobial carryover, and each well was washed three times with sterile deionized water. Biofilms were scraped thoroughly, with a particular attention to well edges (27). The well contents were aspirated and placed in 2 ml of isotonic phosphate buffer (0.15 M, pH 7.2), and the total fungal CFU number was determined by serial dilution and plating on SDA and incubated for 24h at 35°C. Data were log_10_-transformed and were compared with data for untreated biofilms. The MBEC values of drugs were defined as the concentrations of drug that killed 50% (MBEC_50_) or 90% (MBEC_90_) of yeasts in preformed 48-h-old biofilms. All assays included three replicates and were repeated in three independent experiments.

### Cytotoxicity assay

The cytotoxicity of MYC-057 was determined using two human L2 and A549 cell lines (from Melanie Cushion’s laboratory at the University of Cincinnati (Cincinnati, OH)) (59). Briefly, cultured cells were plated at 2 × 10^5^/ml in DMEM/F12 medium supplemented with L-glutamine (2.45 mM), 50 IE/ml penicillin, 50 µg/ml streptomycin and 10% fetal calf serum in a moist atmosphere with 5% (v/v) CO2 and grown at 37°C to conflcent monolayers. The medium was removed and replaced with fresh medium containing the control agent Antimycin A and increasing concentrations of test compounds. Cell viability was evaluated in triplicate assays at three time points (24, 48, and 72 h), performed in three wells. The medium was aspirated from the wells, the adherent cells were lysed with 0.1 M NaOH, and ATP content was assayed in a portion of the lysate using the luciferin-luciferase assay, according to the protocol provided with the ATP-Lite assay kit (Perkin-Elmer, Waltham, MA). The background luminescence was subtracted and replicate well readings were averaged. For each time point, percent reduction in ATP content in all groups and IC50 was calculated as for the in vitro Pneumocystis ATP assays.

### Hemolytic activity

Hemolysis assay was used to determine the potential toxicity of MYC-053. Human erythrocytes from a healthy adult donor were used, as described previously (60,61,62), with a few modifications. MYC-053 was mixed with a 3% (v/v) suspension of washed erythrocytes (in sterile phosphate-buffered saline) for final MYC-053 concentrations of 100–1000 μg/ml, and incubated at 37°C for 3 h in a 96-well plate (Sarstedt). A solution (0.1%) of Triton X-100 (G-Biosciences, St. Louis, MO), a known hemolytic agent, was used as a positive control and DMSO was used as a negative control. Following 6 h of incubation at 36°C, samples were centrifuged and the supernatants were transferred to a 96-well plate to evaluate erythrocyte lysis at 405 nm (Stat fax-2100, Awareness Technology Inc., Palm City, FL) (60).

### Statistical analysis

The Mann-Whitney *U*-test was used to evaluate the differences between antifungal-treated and control samples. Differences at *P* ≤ 0.05 were considered significant. The non-parametric paired Wilcoxon signed-rank test was employed to analyze the pre- and post-challenge differences, and *P* < 0.05 was considered significant. All assays were conduced in triplicate, and were repeated in three independent experiments.

## ACKNOWLEDGMENTS

This work utilized NIAID’s suite of preclinical services for in vitro assessment (Contract No. HHSNHHSN272201100018I, with University of Texas Health Sciences Center, San Antonio). We also like to thank Nathan Wiederhold and Thomas Patterson for performing the MIC testing and for providing additional advices.

## Ethical approval

Not required.

